# Aging induces Nlrp3 inflammasome dependent adipose B cell expansion to impair metabolic homeostasis

**DOI:** 10.1101/607192

**Authors:** Christina D. Camell, Aileen Lee, Patrick Günther, Emily L. Goldberg, Olga Spadaro, Yun-Hee Youm, Andrzej Bartke, Gene B. Hubbard, Yuji Ikeno, Nancy H. Ruddle, Joachim Schultze, Vishwa Deep Dixit

**Author notes:** Address and Correspondence to: Vishwa Deep Dixit, Ph.D., Section of Comparative Medicine and Department of Immunobiology, Yale School of Medicine, 310 Cedar St, New Haven CT 06520.

## Abstract

Visceral adiposity in elderly is associated with alterations in adipose tissue immune cells leading to inflammation and metabolic dysfunction. The Nlrp3 inflammasome is a critical regulator of macrophage activation, inflammation, and immunometabolism in visceral adipose tissue during aging; however, the potential contribution of adipose tissue B cells is unexplored. Here, we show that aging expands adipose-resident B cells and fat-associated lymphoid clusters (FALCs) in visceral white adipose tissue. Adipose tissue B cells exhibit a memory-like B cell profile similar to the phenotype of aged B cells that are increased in spleen of old mice. Mechanistically, the age-induced FALC formation and adipose B cell expansion, but not B cell transcriptional program, is dependent on the Nlrp3 inflammasome. Furthermore, B cell depletion in aged mice restores lipolysis and defense against loss of core body temperature during cold stress. These data reveal that inhibiting Nlrp3-dependent B cell accumulation can be targeted to reverse metabolic impairment in aging adipose tissue.

**Highlights:** - Adipose-resident aged B cells are increased in fat-associated lymphoid clusters (FALC)

- FALC formation and adipose-resident B cell expansion during aging are regulated by the Nlrp3 inflammasome

- Nlrp3 and B cell depletion in aging restores lipolysis and improves cold tolerancea

## Introduction

Increased visceral adiposity that is seen in aged individuals is accompanied by a decreased ability of adipose tissue to maintain homeostatic functions (Kennedy et al., 2014; Stenholm et al., 2008). These core functions of white adipose tissue include lipid storage and endocrine capability, both of which require intricate coordination between adipocytes and resident hematopoietic cells (Hotamisligil, 2006; Kanneganti and Dixit, 2012). White adipose tissue is highly heterogeneous containing defined microenvironment niches in which tissue resident macrophages have distinct roles that facilitate tissue maintenance. Niches such as crown-like-structures (CLS), in which dying adipocytes are cleared by macrophages (Cinti et al., 2005; Martinez-Santibanez and Lumeng, 2014), and sympathetic nerve fibers, in which nerve-associated macrophages regulate local access to catecholamines that stimulate lipolysis (Bartness et al., 2014; Camell et al., 2017; Pirzgalska et al., 2017), have been implicated in metabolic pathogenesis during aging and obesity. Fat-associated lymphoid clusters (FALCs), predominantly composed of B1-innate B cells, serve as unique immunological sites that are acutely responsive to pathogens and increase with chronic inflammation (Benezech et al., 2015; Jackson-Jones et al., 2016; Lumeng et al., 2011; Morris et al., 2013). The contribution of FALCs and FALC-resident cells to age-related metabolic dysregulation remains unknown.

Adipose B cells have many functionally distinct roles, including antibody production, pro- and anti-inflammatory cytokine production and antigen presentation (Benezech et al., 2015; Frasca and Blomberg, 2017; Nishimura et al., 2013; Winer et al., 2011). In diet-induced-obesity, B cells produce IgG antibody and drive Th1 polarization contributing to insulin resistance and clearance of adipocytes in CLS (McDonnell et al., 2012; Winer et al., 2011). However, B1 and B regulatory subtypes favor insulin sensitization via IgM antibodies and anti-inflammatory cytokine production (Nishimura et al., 2013; Shen et al., 2015). Recent work suggests that B2 lymphocytes increase with age in epididymal adipose of mice and ablation of the B cell-specific nuclear cofactor, Oct coactivator from B cells (OcaB), which controls B cell development, improves insulin-sensitivity in middle-aged mice (Carter et al., 2018) and aged B cell (ABC)-like B cells produce TNFα and IgG2c (Frasca and Blomberg, 2017); however, mechanisms that link impaired adipose tissue function to B cell homeostasis in aging are not well understood.

Recent studies demonstrate that the Nlrp3 inflammasome is among the major regulators of age-related inflammation and metabolic disturbance (Bauernfeind et al., 2016; Camell et al., 2017; Spadaro et al., 2016; Youm et al., 2013; Youm et al., 2012). The Nlrp3 inflammasome is an intracellular pattern recognition receptor in innate immune cells that is activated by a wide range of damage associated molecular patterns (DAMPs) (Kanneganti et al., 2007; Schroder and Tschopp, 2010). The identification of the Nlrp3 inflammasome in driving a number of age-related pathologies underscores the importance of innate immune cell-specific inflammation in aging (Bauernfeind et al., 2016; Goldberg and Dixit, 2015; Latz and Duewell, 2018; Youm et al., 2013; Youm et al., 2012).

We have previously shown that aging is associated with reduction in visceral adipose tissue macrophage subsets which lack M1 or M2 polarization, but display senescent-like gene expression signatures that is in part dependent on the Nlrp3 inflammasome (Camell et al., 2017). Here we report that aging is associated with an expansion of adipose-resident B cells in FALCs of white visceral adipose tissue. To determine the nature and function of adipose B cells, we performed flow cytometry phenotyping, whole mount confocal imaging and whole transcriptome expression analysis, which revealed the memory B cell profile. We found that ablation of the Nlrp3 inflammasome in aging protects against adipose tissue-resident B cell expansion and FALC development, without altering the inflammatory transcriptional profile of resident adipose B cells. Thus, we uncovered that age-related expansion of adipose-resident B cells and FALC development requires the Nlrp3 inflammasome-dependent chronic inflammation. B cell depletion in aged mice restored the aberrant adipose immune cell composition and improved metabolic capacity in adipose tissue of aged mice.

## Results

### Aged white visceral adipose shows a unique increase in memory B lymphocytes in FALCs

Since there is an age-related decrease in visceral adipose tissue macrophages (Camell et al., 2017) along with an expansion of visceral adipose tissue CD3^+^ lymphocytes and Tregs (Cipolletta et al., 2015; Lumeng et al., 2011), we sought to investigate the impact of aging on adipose B lymphocytes. To discriminate tissue resident cells from leukocytes in adipose tissue vasculature, circulating cells were labelled with intravenous injection of a CD45.2 antibody (Figure S1A). Mice were sacrificed at 3min post injection, which permitted labeling of all circulating cells, without CD45.2 antibody leaking into tissues or labelling cells extravasating into tissue (Anderson et al., 2014). Adipose tissue digestion was followed by staining with CD45 clone 30-F11, which allows clear separation of tissue resident cells (CD45 clone 30-F11^+^ CD45.2^−^) from circulating cells (CD45 clone 30-F11^+^ CD45.2^+^) (Figure S1A).

Quantification of tissue resident CD3^+^ and B220^+^ cells in mice revealed an increase in CD3^+^ lymphocytes and a 10-fold increase in the percentage of B220^+^ lymphocytes in the visceral adipose from 24 month old wild-type mice (Figure 1A & B). Strikingly, while only 20% of B220^+^ cells adipose tissue from 3 month old mice were tissue resident, nearly 100% of B220^+^ cells from 24 month old adipose tissue were tissue resident (Figure 1C). Furthermore, age-related expansion in adipose tissue resident B cells was observed by 15 months of age and was pronounced in female mice and was less profound, and not statistically significant, in male animals (Figure 1D). Moreover, analysis of tissues from 2 or 24 month old wild-type female mice housed at separate animal facility (Pennington Biomedical, Baton Rouge) also showed a similar increase in visceral adipose B cells (Figure S1B). These results demonstrate that the expansion of resident adipose B cells initiates by 15 months of age and is housing facility-independent, but sex-dependent.

**Figure 1.**
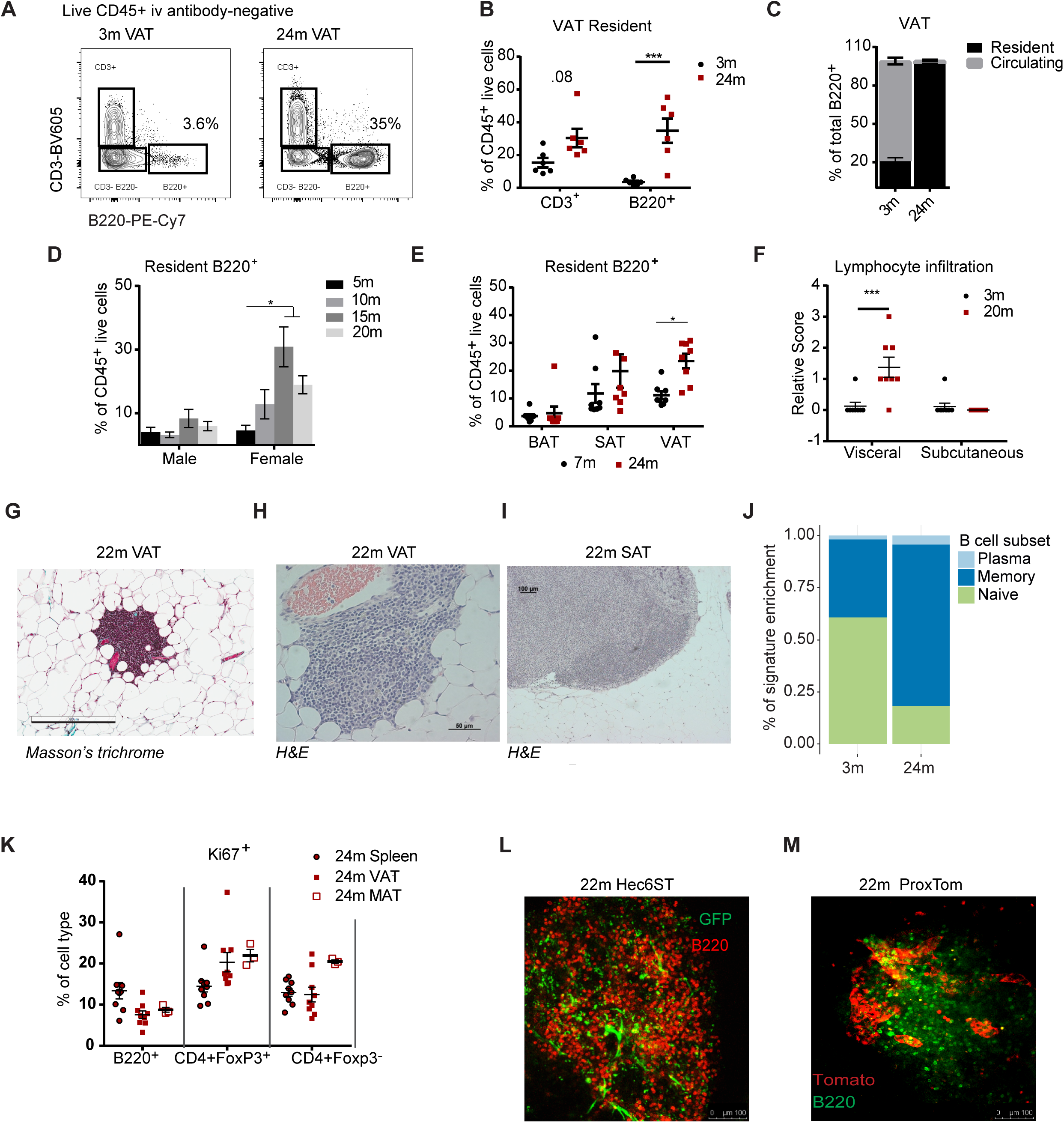
Aging increases FALCs and B cell accumulation in visceral adipose tissue. a. Contour plots gated through CD45^+^ iv antibody^-^ live cells in stroma vascular fraction (SVF) of 3 and 24 month VAT. Values represent the mean percentage of B220^+^ cells. b. Quantification of CD3^+^ and B220^+^ lymphocytes as a percentage of live CD45^+^ residents. Each symbol represents one mouse. c. Quantification of resident and circulating cells as a percentage of total B220^+^ cells from 3 or 24 month old mice. d. Quantification of B220^+^ lymphocytes as a percentage of live CD45^+^ iv^-^ cells, in male or female mice at 5, 10, 15 or 20 months of age. e. Quantification of resident (iv antibody^−^) B220^+^ cells in brown (BAT), subcutaneous (SAT) and visceral adipose tissue (VAT). Each symbol represents one mouse. f. Quantification of lymphocyte infiltration based on hematoxylin and eosin (H&E) staining in the VAT and SAT of 3 or 20 month old mice. Each symbol represents one mouse. g. Masson’s trichrome staining showing a FALC in 22 month old VAT. h. H&E staining showing a FALC in 22 month old VAT. i. H&E staining showing inguinal lymph node in SAT of 22 month old mouse. j. Linear support vector analysis showing gene signature enrichment of VAT B cells that overlap with plasma (light blue), memory (dark blue) or naïve (green) B cells (Newman et al., 2015). k. Quantification of percentage of Ki67^+^ cells, as a proportion of cell subsets, in spleen, VAT or mesenteric adipose tissue (MAT) cells of 24 month old mice. l. Whole mount confocal imaging in 22 month old visceral adipose of Hec6ST mice to visualize HEVs (green) in B220^+^ FALCs (red). m. Whole mount confocal imaging in 22 month old visceral adipose of ProxTom mice to visualize lymphatic vessels (tomato) in B220^+^ FALCs (green). All error bars represent mean±SEM. *<0.05, **<0.01, ***<0.005. Statistical significance was determined by an ANOVA with posthoc test to adjust for multiple corrections.

Excess white visceral adipose tissue is associated with increased risk for metabolic impairments, whereas adipose tissue depots such as subcutaneous white or brown adipose have been linked to insulin sensitivity and thermogenic capacity (Despres and Lemieux, 2006; Zamboni et al., 1997). To determine whether the age-related expansion in resident B cells occurs in the differing adipose depots, we quantified the resident (i.v. CD45.2-FITC negative) B cells from white visceral, white subcutaneous inguinal and brown intrascapular adipose of 7 month or 24 month old mice (Figure S1C). The results show the age-related expansion in resident B cells was specific to white visceral adipose tissue (Figure 1E). In line with quantification using flow cytometry, histopathological scoring analysis revealed significant increases in the lymphocyte infiltration in the visceral, but not subcutaneous adipose tissue of 20 month old mice (Figure 1F). These results suggest an association of adipose B cell expansion with visceral adiposity and the accompanying metabolic risk seen in aging.

Previous reports describe immune cells within white visceral adipose niches influence lipolysis, energy metabolism and tissue functionality (Camell et al., 2017; Xu et al., 2013). To better understand the distribution of lymphocytes throughout two visceral adipose depots, we performed whole mount imaging on an inverted microscope in fixed, but not clarified, mesenteric and gonadal visceral adipose tissues from 7 or 24 month old wild type mice. In 7 month old mice, CD3^+^ and B220^+^ staining was focused in mesenteric lymph nodes of mesenteric visceral adipose and labeling was not seen throughout mesenteric nor gonadal visceral adipose (Figure S1D). Both mesenteric and gonadal adipose tissues from 24 month old mice showed CD3^+^ and B220^+^ staining in clusters, called fat-associated lymphoid clusters (FALCs), which are known to increase with age (Benezech et al., 2015; Lumeng et al., 2011; Moro et al., 2010).

Consistently, histological analysis showed accumulation of lymphocytes within FALCs (Figure 1G&H) that were distinct from lymphocytes seen in a capsule-lined lymph node in subcutaneous inguinal adipose tissue (Figure 1I). Whole mount confocal imaging revealed B220^+^ B and CD3^+^ CD4^+^ T cells in the FALCs from adipose of the 22 month old wild-type mice (Figure S1E). These data demonstrate that aging induces striking increases in resident adipose B cells that are localized in FALCs in mesenteric and gonadal visceral adipose tissues of female mice.

During acute infection B1 cells, an innate-like B cell expressing CD11b and secreting natural IgM antibodies, in FALCs are activated and contribute to pathogen clearance (Jackson-Jones et al., 2016; Moro et al., 2010). To address whether the chronic inflammation seen in aging expands specific populations of B cells, we evaluated the mean fluorescence intensity (MFI) of CD11b and the proportions of CD11b-positive and negative B cell populations within visceral adipose of 3 and 24 month old wild-type male and female mice. CD11b MFI was comparable in the resident visceral adipose B cells from 3 or 24 month old female mice and was not significantly different in males (Figure S1F). Furthermore, there were no differences in the proportions of CD11b^+^ (B1) or CD11b^−^ (B2) resident B220^+^ cells (Figure S1G), suggesting that B cell subsets are equally expanded with age in female mice. To better understand the phenotype of age-expanded visceral adipose B cells, we sorted B220^+^ cells from adipose tissue of 3 or 24 month old wild-type mice for whole transcriptome analysis. Linear support vector regression analysis of sorted B220^+^ cells from visceral adipose tissue revealed that B cells from 24 month old wild-type mice express a predominantly memory-like B cell transcriptional profile (Figure 1J) (Newman et al., 2015). These data demonstrate that the age-related expansion in visceral adipose B cells is primarily an increase in memory-like B cells.

During aging, the accumulation of a unique subset of B cells, termed aged B cells (ABCs), in the spleen, bone marrow and blood has been well described (Hao et al., 2011). ABCs are CD93^−^ CD21^−^ CD23^−^ and evidence suggests that they are antigen-experienced, arising from their encounter with nucleic acid-containing antigens in the context of promoting cytokine production (Naradikian et al., 2016). Consistent with these reports, in 24 month old wild-type mice 20-40% of splenic B cells were CD93^−^ CD21^−^CD23^−^, but nearly 100% of adipose B cells lacked CD93, CD21 or CD23 suggesting that the resident adipose B cells are similar to ABCs found in spleen (Figure S1H). Notably, aged adipose-conditioned media was able to drive the phenotype of CD21^−^ CD23^−^ B cells from 3 month old splenic B cells (Figure S1I). Taken together these results suggest that FALCs act as unique adipose niches during aging with an expansion of resident memory B cells with phenotype specific to white visceral adipose tissue.

### Adipose B cell express Ki67 and have access to lymphatic vessels

Resident adipose tissue immune cells, such as macrophages in response to high-fat diet, (Amano et al., 2014; Zamarron et al., 2017) and CD4^+^ Foxp3^+^ T regulatory cells during aging (Kolodin et al., 2015), are self-maintained through proliferation. To evaluate whether adipose B cells in aged mice are actively dividing, we assessed Ki67 expression, a nuclear antigen expressed in proliferating cells, in B cells and in FoxP3^+^ and FoxP3^-^ CD4^+^ T cells from visceral and mesenteric adipose tissue and spleen of 24 month old wild-type mice. Approximately 10% of B cells from spleen and adipose tissue are positive for Ki67 (Figure 1K). These data indicate that a portion of age-expanded adipose B cells are actively proliferating.

Lymphoid clusters in omentum contain high endothelial venules (HEVs), specialized post-capillary venules essential for naïve lymphocyte trafficking (Buscher et al., 2016; Rangel-Moreno et al., 2009). Whether HEVs and lymphatic vessels are present in the FALCs of aged visceral adipose tissue is unclear. To address whether HEVs and lymphatic vessels are found in FALCs, we imaged FALCs from aged reporter mice specific for HEV or lymphatic vessel markers (Ruddle, 2014; Truman et al., 2012). The 22 month old Hec6ST reporter mice, which express GFP in Hec6ST^+^ cells, showed HEVs within FALCs and in close proximity to B220^+^ clusters (Figure 1L). Likewise, 22 month old ProxTom mice, which express intracellular tomato in the Prox1^+^ lymphatic endothelial cells, showed lymphatic vessels within FALCs (Figure 1M). Interestingly, we also found lymphatic vessels in adipose tissue that were not within FALCs, but that still contained B cells (Figure S1J). To confirm the presence of Prox1^+^ lymphatic vessels in wild-type aged mice, we optimized the nuclear antibody staining protocol for Prox1 on whole mount mesenteric lymph nodes as a positive control (Figure S1K). Fluorescence analysis of whole mount adipose tissue revealed Prox1 staining surrounding FALCs even at low magnification (Figure SlL). Furthermore, confocal microscopy revealed clear Prox1+ nuclear staining in vessels near and surrounding FALCs (Figure S1M). These data demonstrate that HEVs and lymphatic vessels are found in FALCs in aged mice suggesting that FALCs support lymphocyte trafficking.

Taken together these experiments identified a dramatic increase in the resident population of B lymphocytes in FALCs of white visceral adipose tissue from female mice. Adipose B cells display transcriptional signatures akin to memory B cells and exhibit the characteristics of ABCs. Our data also suggest that in aged mice adipose-resident B cell expansion may be supported by both active proliferation and local trafficking via lymphatic vessels and HEVs.

### Age-induced Adipose B cell expansion and FALC formation in adipose is regulated by the Nlrp3 inflammasome

In the context of acute inflammation, macrophage-derived TNFα promotes stromal cell activation and the accumulation of cells in FALCs (Benezech et al., 2015). Given the association of adiposity with chronic inflammation during aging, we wished to explore the spatial association of macrophages with FALC B cells. To track myeloid cells in FALCs, we aged and imaged mT/mG;LysM-cre reporter mice, which allowed for simultaneous visualization of myeloid cells (GFP^+^) and all cells (tomato^+^). These analyses showed dense capillary networks within FALCs indicative of the vascular network as previously reported (Benezech et al., 2015). Antibody staining showed FALCs contained numerous B220^+^ cells, some of which could be found in close contact with macrophages (Figure 2A; S2A). Given this close association of macrophages to B cells within FALCs and our previous findings showing macrophage and Nlrp3 inflammasome regulation of adipose tissue homeostasis (Camell et al., 2017; Vandanmagsar et al., 2011), we asked whether the age-related expansion in adipose B cells requires the Nlrp3 inflammasome. Compared to 24 month old wild-type female mice, Nlrp3-deficient mice showed a significant reduction in the percentage and the total numbers of B cells in visceral adipose (Figure 2B). Interestingly, although CD4^+^ and CD8^+^ T cells are increased with age, there were no statistically significant differences when comparing the 24 month old wild-type and Nlrp3-deficient mice (Figure S2B), indicating that the Nlrp3 inflammasome specifically promotes the age-related B cell expansion in visceral adipose tissue at 24 months of age, without affecting age-related T cell expansion.

**Figure 2.**
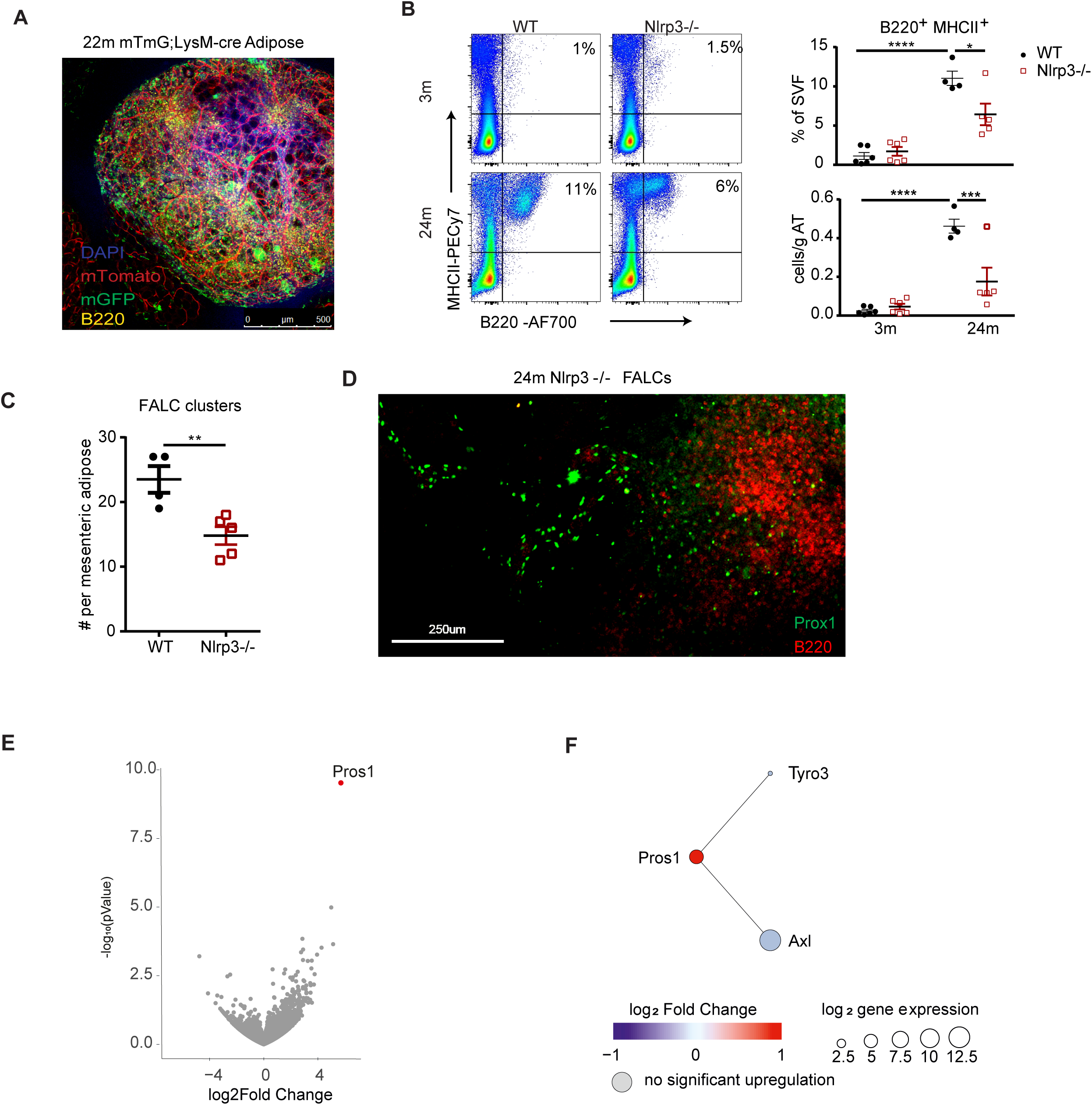
Nlrp3 inflammasomes controls B cell expansion and FALC development. a. Whole mount confocal imaging of FALC in VAT of 22 month old mT/mG;LysMcre with B220 (yellow) and DAPI (blue) antibody staining. mTomato expressed on all cells and mGFP on myeloid cells. b. Flow cytometry plots showing B220^+^ MHCII^+^ cells from the visceral adipose tissue of 3 or 24 month wild-type and Nlrp3-/- mice. Quantification of the percentage and cells (x10^6)/g tissue of B220^+^ MHCII^+^ cells. c. Quantification of the numbers of FALCs per adipose tissue in 24 month old wild-type or Nlrp3-deficient adipose tissue. d. Whole mount confocal imaging to visualize Prox1^+^ (green) lymphatic endothelial cells in FALCs (B220^+^; red) e. Differentially expressed genes in visceral adipose B cells from 24 month old wild-type and Nlrp3-/- mice. Grey dots represent no significant change in gene expression, red dots represent significant changes in gene expression (FDR<0.1). f. Schematic visualizing Pros1 change in WT and Nlrp3-/- visceral adipose B cells and potential interactions with TAM receptors on macrophages All error bars represent mean±SEM. *<0.05, **<0.01, ***<0.005. Statistical significance was determined by an ANOVA with posthoc test to adjust for multiple corrections.

To determine whether the Nlrp3 inflammasome is required for visceral adipose T cell expansion at an earlier age, we performed flow cytometric quantification in young wild-type mice (4 month old) and middle-aged (10 months old) wild-type or Nlrp3-deficient mice, a time at which B cell expansion has not yet occurred. Consistent with our previous findings in 24 month old mice (Camell et al., 2017), the 10 month old wild-type mice showed reduced CD11b+ myeloid cells, which was prevented by Nlrp3-deficiency (Figure S2C). Furthermore, adipose tissue CD4^+^ T cells were significantly elevated in the 10 month old wild-type mice, and prevented by Nlrp3-deficiency (Figure S2D). As expected, there was no age-related expansion in visceral adipose B cells in the wild-type mice at 10 months of age (Figure S2E). These data support a model in which the Nlrp3 inflammasome initially controls age-related increases in total CD4^+^ T cells, and by 24 months of age, the Nlrp3 inflammasome is specifically required for resident adipose B cell expansion.

We next asked whether the Nlrp3 inflammasome is also required for FALC development. Compared to 24 month old wild-type mice, there was a significant reduction in the number of FALCs per adipose tissue in the aged Nlrp3-deficient mice (Figure 2C). To determine whether the reduced FALC formation is due to a reduction in lymphatic vessels found in FALCs, we performed whole mount confocal imaging for Prox1^+^ lymphatic endothelial cells in the adipose tissue of the 24 month old wild-type and Nlrp3-deficient mice. All FALCs in adipose tissue contained Prox1^+^ lymphatic vessels (Figure 2D). These data suggested that although the Nlrp3 inflammasome is required for age-related B cell expansion, lymphatic vessel formation is Nlrp3 inflammasome-independent.

Given that Nlrp3-deficiency in aging protects against adipose B cell expansion, we next investigated the inflammatory potential of these age-expanded FALC B cells using unbiased RNA sequencing analyses. Surprisingly, this analysis revealed that compared to 24 month old wild-type mice, the adipose B cells from age-matched Nlrp3-deficient mice displayed almost no differentially expressed genes. Only a single gene, Pros1, was identified to be significantly regulated in the adipose tissue B cells from Nlrp3-deficient animals. Pros1, which encodes Protein S, is significantly elevated in the adipose B cells from Nlrp3-deficient mice (Figure 2E), and interacts with Tyro3, Axl and Mer (TAM) receptors (Rothlin et al., 2015). Pros1-TAM receptor interaction has potential to drive anti-inflammatory signaling in macrophages (Figure 2F). There were no other alterations in pro- or anti-inflammatory genes that were differentially regulated by Nlrp3-deficiency. Taken together with our previous findings (Camell et al., 2017), these data describe an Nlrp3 inflammasome-dependent role for age-related changes in adipose tissue resident macrophages and B cells. In contrast to the Nlrp3-dependent transcriptional regulation identified in macrophages (Camell et al., 2017), the transcriptional profile in B cells is primarily regulated in an Nlrp3-independent manner. These data indicate that Nlrp3 inflammasome-derived inflammation from macrophages acts upstream of visceral adipose B cell expansion and controls resident B cell accumulation without altering their transcriptional phenotype. Thus, the Nlrp3 inflammasome-mediated reduction in adipose tissue inflammation during aging may result in part from numerical reduction in B cells.

### Nlrp3-driven macrophage interactions do not drive FALC development

Macrophage-mediated inflammation is required for FALC development by directing stromal cell activation to initiate lymphocyte recruitment (Benezech et al., 2015). Given the role of the Nlrp3 inflammasome in resident B cell and FALC accumulation in aged adipose tissue, we next explored the possibility that aged adipose tissue macrophages directly control B cell recruitment. Using our previous adipose macrophage whole genome sequencing analysis, (GSE93202) (Camell et al., 2017), we examined the mean chemokine gene expression in wild-type or Nlrp3-deficient adipose macrophages from 3 or 24 month old mice. Surprisingly, hierarchical clustering analysis of chemokine expression did not reveal the group structure within the dataset suggesting that the Nlrp3 inflammasome is not a predominant regulator of B cell trafficking in aging adipose tissue (Figure S2F). To address the possibility that macrophage-B cell interaction contributes to the Nlrp3 inflammasome-regulated resident B cell expansion, we generated a macrophage-B cell interaction model by first curating a list of all possible receptors and ligands using the Fantom5 interaction database. To focus on the Nlrp3-dependent contribution, we identified macrophage ligands and receptors which are regulated in an Nlrp3 inflammasome-dependent manner and which have corresponding receptors expressed in the adipose tissue B cells (Figure S2G). All macrophage ligands and receptors identified showed potential to interact with B cells in one or more ways. Some interactions implied inflammatory signaling regulation (eg. Tlr4-S100a8/9 and Tnfrsf18-Tnfsf18 interaction) between macrophages and B cells, but no single interaction appears sufficient to explain the development of FALCs, suggesting that in context of low-grade chronic age-induced inflammation multiple interactions contribute to adipose-resident B cell expansion during aging.

### Reduced Nlrp3 driven inflammation and longevity is linked to lower ABC accumulation

The increase in an antigen-experienced subset of B cells, called aged B cells (ABCs) in the spleen of aged mice has been attributed to chronic levels of toll-like receptor and B cell receptor signaling (Hao et al., 2011; Rubtsova et al., 2015; Russell Knode et al., 2017). Given the similarity of adipose B cells to ABCs (Figure S1H), the ability of adipose to drive ABC formation (Figure S1I) and the dependency on the Nlrp3 inflammasome for adipose B cell expansion (Figure 2B), we wondered whether the age-related increase in splenic ABCs requires Nlrp3 inflammasome. Quantification of CD21^−^CD23^−^ B cells in the spleen of 3 month or 24 month old wild-type and Nlrp3-deficient mice revealed significant elevation in the frequency of ABCs in the 24 month old wild-type mice that was less pronounced in the 24 month old Nlrp3-deficient mice (Figure 3A). We identified a corresponding age-regulated decrease in marginal zone and follicular B cells that was not restored with Nlrp3-deficiency (Figure 3B), suggesting that the Nlrp3 inflammasome is required for age-related ABC expansion, but not marginal or follicular B cell alterations.

**Figure 3.**
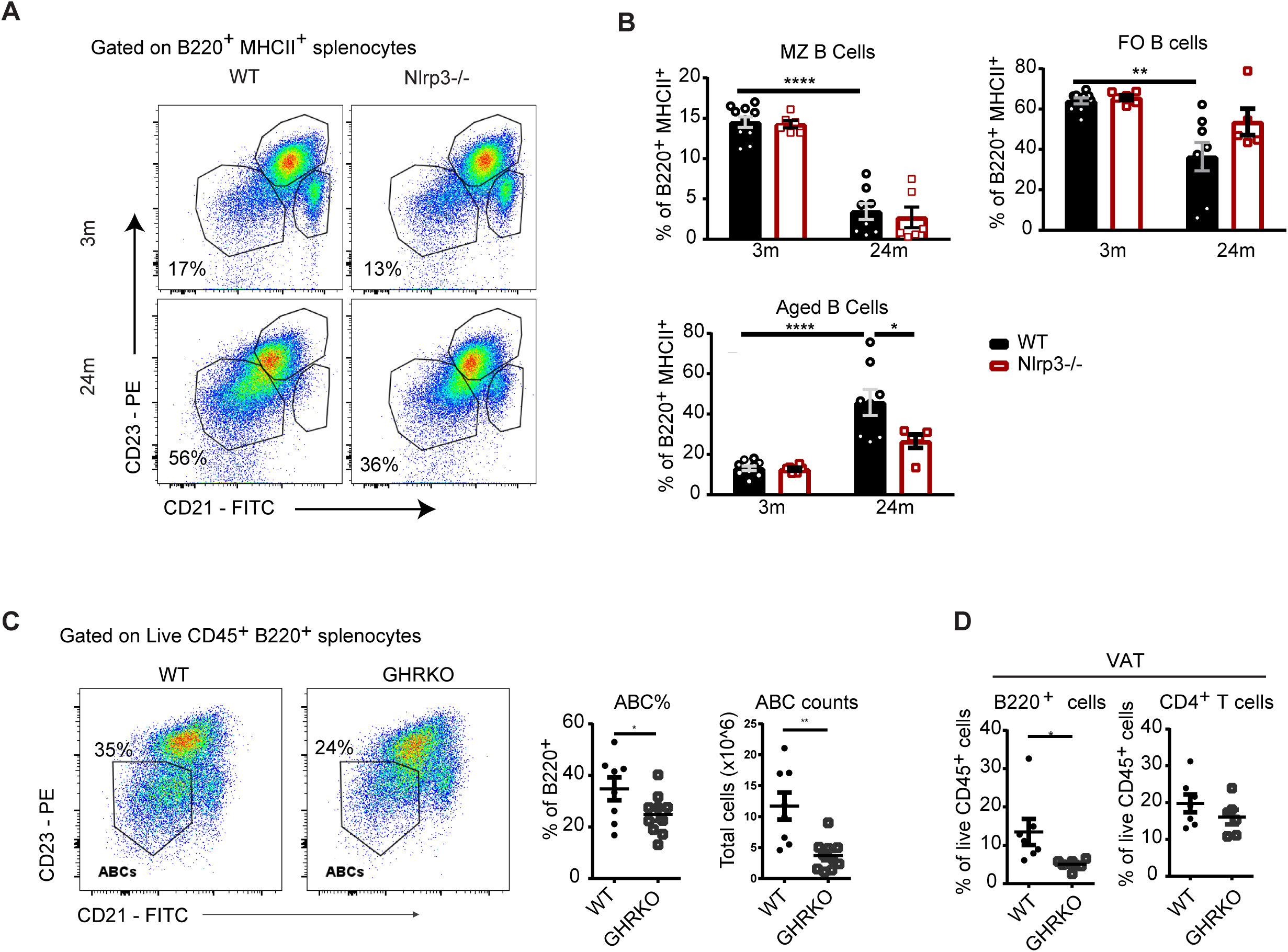
B cell expansion is associated with reduced healthspan and lifespan. a. Pseudocolor density plot gated through CD45^+^ B220^+^ MHCII^+^ splenocytes to show B cell subsets in 3 or 24 month old WT and Nlrp3-deficient mice. b. Quantification of splenic B cell subset frequency. MZ: marginal zone B cells; FO: follicular B cells. c. Representative density plots and quantification and of B cell subsets in B220^+^MHCII^+^ splenocytes from WT or GHRKO mice at 20 months of age. e. Quantification of B220^+^ and CD4^+^ cells from the visceral adipose tissue of WT or GHRKO mice at 20 months of age. All error bars represent mean±SEM. *<0.05, **<0.01, ***<0.005. Statistical significance was determine by an ANOVA with posthoc test to adjust for multiple corrections.

### Long lived-GHRKO mice have reduced adipose cellularity and adipose B cells

Long-lived mice with reduced growth hormone (GH) signaling have improved insulin sensitivity and reduced Nlrp3 inflammasome activation (Spadaro et al., 2016). To address whether longevity is associated with ABCs and visceral adipose B cell expansion in aging, we aged growth hormone receptor (GHR)-sufficient and GHR-knockout (GHRKO) mice to 20 months of age. Along with their extended lifespan and increased insulin sensitivity (Coschigano et al., 2003; Masternak and Bartke, 2012), GHRKO mice have lower body weight, reduced visceral adipose tissue weight and decreased adipose cellularity (Figure S3A). Quantification of ABCs in wild-type or GHRKO spleens revealed significant decreases in the percentage and total number of ABCs (Figure 3C). Furthermore, the visceral adipose tissue of GHRKO mice had significantly reduced percentage of B220^+^ cells, but not CD4^+^ T cells (Figure 3D). These data highlight that reduction of ABCs is associated with extended lifespan and may be related to improved metabolic outcomes associated with longevity.

### Depletion of adipose B cells restores insulin sensitivity of aged mice

As the age-related expansion in adipose B cells is localized to visceral adipose and not to other adipose tissues depots (Figure 1D), we first wanted to determine whether a reduction in visceral adipose tissue B cell numbers would restore age-related insulin resistance. We devised an intra-adipose injection protocol allowing for adipose tissue-specific depletion of adipose B cells (Figure S4A) to avoid systemic depletion of B cells. Body-weight and visceral adipose tissue weight in ISO-treated and mCD20 mAb-treated mice were comparable (figure S4B&C). At 30 days after intra-adipose injection with CD20 mAb, B cells were specifically reduced in visceral, but not in subcutaneous adipose tissue, spleen, bone marrow or peritoneal cavity (Figure 4A & 4C, S4D). When challenged with insulin, 20 month old wild-type mice given the intra-adipose injection with CD20 mAb showed improved insulin tolerance as compared to the 20 month old wild-type mice given an ISO control injection (Figure 4C). As improvements in insulin sensitivity suggested that lipolytic capacity of aged-adipose may also be improved, we fasted the intra-adipose injected mice to induce lipolysis, a process that can be measured by release of glycerol from adipose tissue (Schweiger et al., 2014). However, there were no differences in glycerol release from the fed (Figure 4D) or fasted (Figure 4E) visceral adipose tissue in ISO-treated or CD20 mAb-treated mice. Collectively, these data indicate that specific depletion of the visceral adipose tissue B cells was capable of improving age-related impaired insulin sensitivity, but not lipolysis resistance.

**Figure 4.**
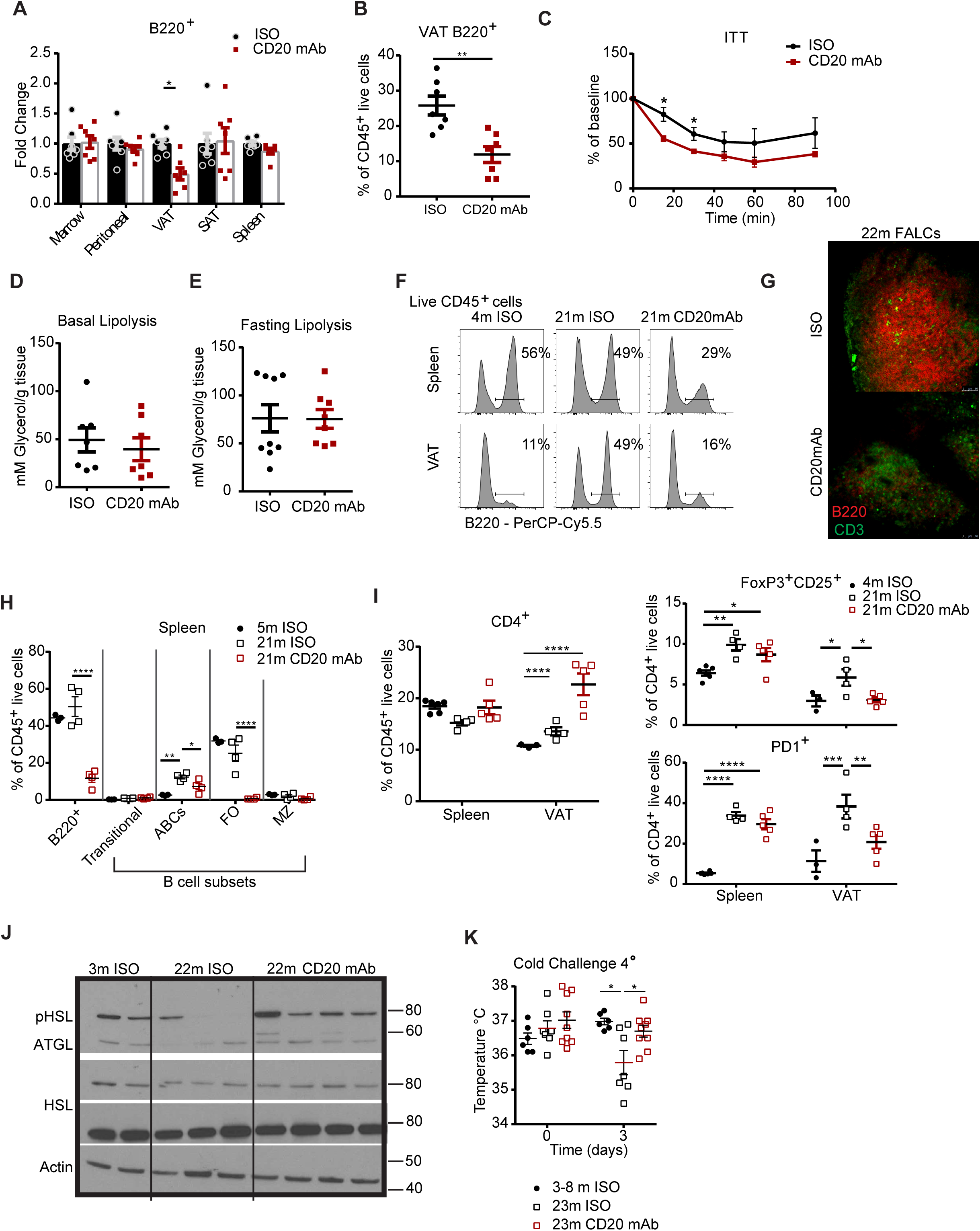
Depletion of B cells restores adipose metabolic capacity. a. Quantification of the fold change (% of B cell from CD20 mAb tissue/ average % of B cell from ISO tissue). Each symbol represents an individual mouse. b. Quantification of the frequency of B220^+^ cells gated through live CD45^+^ cells in visceral adipose of 20 month old wild-type mice given a single intra-adipose injection of ISO or CD20 mAb. Each symbol represents an individual mouse. c. Insulin tolerance test (ITT) in 21 month old wild-type mice treated as indicated. d. Glycerol per gram of tissue in visceral adipose tissue from fed mice given visceral adipose treatment as described. Each symbol represents an individual mouse. e. Glycerol per gram of tissue in visceral adipose tissue from mice that were fasted for 24 hours. Each symbol represents an individual mouse. f. Histogram plots showing B220^+^ cells in tissues from 4 or 21 month old mice given intraperitoneal injection of ISO or CD20 mAb. Values represent the mean percentage for that condition. g. Whole mount imaging showing representative FALCs in mesenteric adipose from 22 month old mice. (Top) FALCs from adipose of mice treated with ISO control, (bottom) FALCs from mice treated with CD20 mAb. h. Quantification of ABCs, FO B cells and MZ B cells in spleen of 5 or 21 month old mice given ISO control or 21 month mice treated with CD20 mAb. i. Quantification of the percentage of CD4^+^ T cells out of the total CD45^+^ live cells in spleen or visceral adipose. (right top) Quantification of CD4^+^ Foxp3^+^ Cd25^+^ cells out of the total CD4^+^ T cells. (right bottom) Quantification PD1^+^ cells out of total CD4+ cells in the spleen and visceral adipose tissue of conditions described. Each symbol represents one mouse. j. Western blot showing phosphorylated HSL, ATGL, total HSL and ACTIN levels in VAT of 3 month, 22 month give isotype control or CD20 mAb as described. Each lane represents one mouse. Representative of two individual experiments. k. Rectal temperature at day 0 and day 3 in wild-type mice treated as described and exposed to four degrees. Each symbol represents one sample. All error bars represent mean±SEM. *<0.05, **<0.01, ***<0.005. Statistical significance was determined by an ANOVA with posthoc test to adjust for multiple corrections.

### Inhibition of systemic B cell expansion restores metabolic homeostasis and thermogenesis in the aged

To further examine the contribution of both age-expanded ABCs in lymphoid tissues and adipose B cells in metabolic dysregulation, we performed intraperitoneal (i.p.) injection of the ISO or CD20 mAb into 21month old mice to systemically deplete all B cells (Figure S4E). As expected, ip injection of CD20 mAb reduced the percentage of B cells in the spleen and visceral adipose tissue in the 21month old mice (Figure 4F). FALCs tended to appear smaller and show reduced staining for B220 (Figure 4G) and percentages of ABCs in the spleen were significantly reduced with CD20 mAb treatment (Figure 4H).

To address whether B cell depletion could restore age-related alterations in immune cell composition, we quantified the percentage of total CD4^+^ T cells, as well as, the PD1^+^ effector population and Treg cells. There was an increase in the frequency of total CD4^+^ cells (Figure 4I) in the B cell depleted mice, but there was a statistically significant visceral adipose-specific reduction in the Treg and CD4^+^PD1^+^ sub-populations (Figure 4I, S4H and S4I). These data indicate that although a single CD20 mAb injection to deplete B cells does not reduce CD4^+^ T cell frequency, the CD4^+^ T cell subsets are more similar to subsets of CD4^+^ T cells from young adipose.

Consistent with previous findings showing restored insulin tolerance in obese mice depleted of B cells and in aged mice depleted of Tregs (Bapat et al., 2015; Winer et al., 2011), 20 month old B cell-depleted mice had improved insulin sensitivity (Figure S4J and S4K). To address whether systemic depletion of B cells could restore lipolytic signaling and metabolic substrate availability in aged mice, 22 month old mice given an intraperitoneal injection of ISO antibody or CD20 mAb were fasted for 24hours. The adipocyte lipases, adipose triglyceride lipase (ATGL) and hormone sensitive lipase (HSL) are required for triglyceride hydrolysis during fasting and are reduced in fasted aged visceral adipose tissue (Camell et al., 2017). As compared to the ISO treated aged wild-type mice, CD20mAB treatment resulted in increased levels of phosphorylated HSL and total ATGL during fasting, indicating that the age-related reduction in lipolytic signaling was restored with B cell depletion (Figure 4J). Consistent with improved lipolytic capability, the visceral adipose tissue from mice given CD20mAB also had increased isopreteronol-stimulated lipolysis (Figure S4L). Given that white adipose lipolysis is required for the generation of metabolic substrates to induce thermogenesis and protect against cold stress (Schreiber et al., 2017; Shin et al., 2017) and aged mice have impaired maintenance of core body temperature during cold stress (Figure S4M) (Berry et al., 2017), we wanted to test whether B cell depletion would improve age-induced defense against cold stress. Consistent with the restored lipolysis providing additional metabolic substrates, 20 month old B cell-depleted mice were able to maintain core body temperature when placed at four degrees C (Figure 4K). Taken together these data implicate the Nlrp3 inflammasome-induced inflammation and B cell expansion in as mechanisms that contribute towards impaired lipolytic capacity of visceral adipose tissue and inability to defend against cold stress in aging.

## Discussion

In this study we address the mechanism of age-related inflammation and adipose dysfunction by investigating the resident adipose tissue B cell phenotype and function in visceral adipose tissue, an organ implicated in the regulation of lifespan and healthspan. Aging leads to expansion of resident adipose B cells localized to FALCs in white visceral, but not subcutaneous or brown adipose. Reminiscent of the ABC expansion, adipose B cell expansion is primarily evident in female mice. B cells exhibit a memory phenotype and their expansion may require both proliferation and local trafficking. The Nlrp3 inflammasome is required for adipose B cell and FALC expansion in aging, but surprisingly is not required for the inflammatory transcriptional phenotype of resident adipose B cells. Functionally, treatment of aged mice with CD20 mAb caused a reduction in FALC-resident B cells and aged B cells, along with a restoration of CD4^+^ T cell subsets, restoration of lipolysis and improved capability to maintain cold-defense. Our data provide new conclusive information that the Nlrp3 inflammasome drives resident adipose B cell expansion and impaired adipose metabolic capacity during aging.

That inflammation and metabolism are intimately linked, particularly in obesity, is well accepted (Ferrante, 2013; Hotamisligil, 2006). However, how aging disturbs immunometabolic inputs that impact maintenance of adipose tissue and metabolic homeostasis is less clear. Although we have uncovered that the Nlrp3 inflammasome-expansion of resident adipose B cells contributes to lipolysis resistance and CD4^+^ T cell subpopulation expansion, how and why B cells localize to FALCs are likely regulated by additional mechanisms apart from the inflammasome. Additionally, the mechanisms responsible for B cell recruitment to FALCs are particularly intriguing given that B cells do not express the Nlrp3 inflammasome and this process warrants further study.

In the young, the adipose tertiary lymphoid structures are local sites of immune surveillance, contributing to the clearance of TLR agonists and pathogens (Benezech et al., 2015; Jackson-Jones et al., 2016; Moro et al., 2010); however, our data suggest that FALCs also control metabolic capacity of adipose tissue. Lipolysis is critical in infections and specific metabolic substrates help drive an efficient and effective immune response (Gazos-Lopes et al., 2017; Wang et al., 2018). Whether B cells regulate host energy homeostasis during various pathophysiological situations requiring metabolic substrates remains to be determined. Moreover, additional studies are needed to identify whether B cells express circulating factors or act directly to impair lipolytic capacity of adipocytes.

Our data offers novel insight into time-dependent and sex-dependent effects on adipose hematopoietic cells. Female aging is accompanied by loss of hormone production, systemic alterations in immune cell responses, and increased risk for autoimmunity (Ghosh et al., 2014). Given that visceral adipose B cell expansion occurs between 10 and 15 months, which is prior to splenic ABC expansion, but follows decreases in adipose tissue macrophages, CD4^+^ T cell expansion and ovarian failure in female mice (Diaz Brinton, 2012; Gosden et al., 1983; Hao et al., 2011), it is interesting to speculate on potential multiple causal factors of these associated events. Our B cell depletion experiments suggest a direct link between B cell expansion and T cell subset accumulation (Figure 4I). Endogenous sex hormones influence both adaptive and innate immunity and a direct link with the Nlrp3 inflammasome has not been reported.

Lymphatic vessels maintain fluid balance and regulate transport of antigen and immune cells (Ruddle, 2016); however they have also been implicated in driving adipogenesis, adiposity and insulin resistance (Harvey et al., 2005). Ours study identified HEVs and lymphatic vessels within FALCs, suggesting regulated trafficking of lymphocytes. However, given previous findings describing the loss of functionality in lymphatic vessels and increased permeability in aging (Zolla et al., 2015), lymphatic vessels may also feed-forward into chronic age-related inflammation. FALC numbers are reduced in the aged Nlrp3-deficient mice, yet still contain lymphatic vessels; whether improved functionality of lymphatic vessels contributes to improved metabolic capacity remains to be tested.

Together, these data show a role for B cell depletion in restoring adipose tissue immune cell compartments, as well as metabolic and immunologic capacity. Importantly, our data reveal previously unrecognized role of the Nlrp3 inflammasome in control of B cell expansion in aging and raise the possibility that FDA approved monoclonal antibodies to deplete B cells may be repurposed for reducing inflammation and reversing metabolic impairment in the elderly.

## Limitations of the study

This study provides evidence that macrophage expressed Nlrp3 inflammasome regulates B cell homeostasis in aging adipose. B cells do not express Nlrp3 inflammasome and are likely impacted by the inflammasome dependent cytokines IL-1β and IL-18 which increase with age. Since the next steps to test the causal role of one of these cytokines would necessitate aging of cell type-specific IL-1 and IL-18 deficient mice our study could not test the contribution of macrophage derived secreted factors to B cell and FALC formation in aging adipose. The RNA sequencing of B cells from aged control and Nlrp3 deficient mice revealed only upregulation of Pros1, which is known to exert anti-inflammatory signaling via TAM receptors. However, whether induction of Pros1 in aged adipose B cells is sufficient for conferring a protective immunometabolic response is not addressed in this study and requires creation of aging cohorts of cell specific Pros1 deficient mice. The reversal of lipolysis resistance in aged mice post B cell depletion suggest that secreted factors from B cells either impact the catecholamine signaling in adipocytes or the sympathetic nervous system innervation itself. Nonetheless, this study provides new evidence that Nlrp3 inflammasome is a surprising new regulator of adipose B cell expansion in aging and opens new avenues for future investigation of immunometabolic checkpoints of aging.

## Author contributions

C.D.C. carried out most experiments. A.L., E.L.G, O.S., and Y.H. assisted with experiments. Y.I. and G.B.H. performed pathological analysis. A.B. generated the GHRKO mouse model. P.G. and J.S. performed the bioinformatics analysis and interpretation. C.D.C and V.D.D. conceived the project, analyzed the data and wrote the manuscript with input from all co-authors.

## Acknowledgements

We thank Genentech Inc for providing the *Nlrp3*-deficient mice and the CD20mAb. We also thank The Yale Center on Genomic Analysis (YCGA) for RNAseq studies and the P. Cresswell laboratory for confocal microscopy support. J.L.S. was funded by the German Research Foundation (SFB704, SFB645) and by the ImmunoSensation Cluster of Excellence Bonn. C.D.C. was supported by AFAR (American Federation of Aging Research Postdoctoral transition fellowship) and NIA (K99AG058800). E.L.G was supported by AFAR (Postdoctoral fellowship) and NIA (K99AG058800). The Dixit laboratory is supported in part by NIH grants P01AG051459, AI105097, AG051459, AR070811, the Glenn Foundation on Aging Research and Cure Alzheimer’s Fund. A.B. was supported by R01AG019899. AB thanks J.J. Kopchick for providing GHRKO breeders.

## Declaration of Interests

The authors declare no conflict of financial interest.

## STAR METHODS

## CONTACT FOR REAGENT AND RESOURCE SHARING

Further information and requests for resources and reagents should be directed to and will be fulfilled by the Lead Contact, Dr. Vishwa Deep Dixit.

## EXPERIMENTAL MODEL AND SUBJECT DETAILS

### Animal care

All mice were housed in specific pathogen-free facilities in ventilated cage racks that deliver HEPA-filtered air to each cage with free access to sterile water through a Hydropac system at Yale School of Medicine. Sentinel mice in our animal rooms were negative for currently tested standard mouse pathogens (Ectromelia, EDIM, LCMV, *Mycoplasma pulmonis*, MHV, MNV, MPV, MVM, PVM, REO3, TMEV and Sendai virus) at various times while the studies were performed (Research Animal Diagnostic Laboratory). C57BL6/J (wild-type) mice were bred from our colony, purchased from Jackson Laboratories or received from the National Institute of Aging Rodent colony. *Nlrp3*^*-/-*^ have been previously described and were bred in our facility, along with their wild-type controls (Mariathasan et al., 2006). *LysM-cre mT/mG* mice were bred in our facility. *ProxTom* and *HEC6ST-GFP* reporter mice were generated and bred by Nancy Ruddle (Bentley et al., 2011; Truman et al., 2012). GHRKO mice and their controls were generated and bred by Andrzej Bartke (Coschigano et al., 2003). All knockout mice were compared to wild-type controls raised in the same facility. Mice were fed a standard vivarium chow (Harlan 2018S) and housed under 12hr light/dark cycles. Mice were examined and only the mice that did not have lymphoma were used for experiments. All experiments and animal use were conducted in compliance with the National Institute of Health Guide for the Care and Use of Laboratory Animals and were approved by the Institutional Animal Care and Use Committee at Yale University. All mouse models used and the sex and number of animals are listed in the below sections.

### Mouse models

#### For insulin tolerance test

Mice were fasted for 4 hours prior to ITT. Insulin was given by i.p. injection (0.8U/kg) and blood glucose levels were measured by glucometer.

#### For fasting

At 8am, mice were placed into a clean cage without food, but with water ad lib. Fast was complete after 24 hours when mice were euthanized for tissue harvest.

#### For cold challenge

Rectal temperature was taken using a rectal thermometer at 7am each day. Baseline body-temperature was taken at 7am at room temperature housing. Mice were then transferred to a cold room that is maintained at 4°C for three days. 12hour light/dark cycle and food and water were maintained throughout the cold challenge.

#### For B cell depletion

intra-adipose depletion was performed during survival surgery under anesthesia. A small ventral midline skin incision was made to access gonadal adipose tissue. 100ul of isotype control antibody (clone C1.18.4; bio X cell in vivo; catalogue#: BP0085) or CD20 monoclonal antibody (Genetech; clone 5D2) is injected into both left and right adipose tissue depots (50ug/depot). The tissue is replaced into the abdomen and the abdominal muscle and skin will be sutured with 4-0 Vicryl. Mice are monitored daily until incision heals. Systemic depletion was performed by i.p. injection of isotype control or CD20 mAB (100ug).

### Animal Experiments

*Experiment 1*: Young vs old adipose tissue immune cell quantification (two individual cohorts compiled) Wild-type; N=3 at 3m; N=3 at 24m in each individual cohort; Female mice

*Experiment 2*: Time-course and sex comparison to quantify resident adipose B cells Wild-type; N= 5 at each time point for each sex.

*Experiment 3*: Aging of mice in separate animal facility (Pennington Biomedical research center) Wild-type; N=3 at each age; female mice

*Experiment 4*: Adipose depot comparisons of resident adipose B cells in 7m and 24m old female mice Wild-type; N= 8 at 7m; N=7 at 24m; each pooled from multiple litters that were bred and housed at Yale University; Female mice

*Experiment 5*: Quantification of visceral and subcutaneous lymphocyte infiltration Wild-type; N= 8 at 3m and N =8 at 20m; Female mice

*Experiment 6*: Sorting of adipose tissue B cells for RNA sequencing analysis Wild-type; N= 3 at 3m and N =2 at 24m pooled from 2 mice each*

*Experiment 7*: Ki67 expression in immune cells from aged animals (two individual cohorts compiled) Wild-type; N=3 and N=4 at 24m in two individual cohorts

*Experiment 8*: Aging of reporter mice and wild-type mice for confocal analysis Hec6ST-GFP mice N=3; ProxTom mice N=3; wild-type N=3; all 20-22m; female mice

*Experiment 9*: Aging of WT and Nlrp3-/- mice for adipose B and T cell quantification at 3m or 24m of age N = 3 WT at 3m; N= 5 Nlrp3-/- at 3m; N=7 WT at 24m; N=5 Nlrp3-/- at 24m pooled from two separate experiments

*Experiment 10*: Aging of WT and Nlrp3-/- mice for FALC quantification and Prox1+ staining N= 5/5 WT/Nlrp3-/- at 24m for FALC quantification. N= 3/3 WT/Nlrp3-/- at 24m; female mice

*Experiment 11*: Aging of WT and Nlrp3-/- mice for adipose B cell sorting N=2 WT at 24m of age and N=3 Nlrp3-/- at 24m of age pooled from 2 mice each*

*Experiment: 12*: Aging of WT and Nlrp3-/- mice for quantification of adipose tissue B and T cells at 4m and 10m of age. N= 4; female mice at 4m WT, 10m WT and 10m Nlrp3-/-

*Experiment 13*: Aging of WT and GHRKO mice for immune cell quantification N = 7 WT and N=9 GHRKO at 20m, pooled from two separate experiments

*Experiment 14*: Intra-adipose injection of CD20 mAB for visceral adipose-specific B cell depletion (two individual cohorts) N=7 ISO and N=7 CD20mAB; pooled from two separate experiments; female mice

*Experiment 15*: I.P. injection of CD20 mAB for systemic depletion of B cells for insulin tolerance test N = 4 ISO and N= 5 CD20 mAB

*Experiment 16*: I.P. injection of CD20 mAB for systemic depletion of B cells with fasting challenge (two individual cohorts) N= 3 3m ISO; N= 3 22m ISO; N= 4 22m CD20 mAB; repeated twice; representative data shwon

*Experiment 17*: I.P injection of CD20 mAB for systemic depletion of B cells with cold challenge (three individual cohorts compiled) N= 6 3-8m ISO; N= 7 23m ISO; N=9 23m CD20m AB pooled together for rectal temperature

All N’s represent a single animal (biological replicate), except for RNA sequencing experiments which required pooling of adipose tissue from mice to collect the N’s indicated.

## METHOD DETAILS

### Adipose digestion and stromavascular staining

Visceral adipose was collected after euthanization and weighed. Tissue was enzymatically digested in 0.1% collagenase I (Worthington Biochemicals) in HBSS (Life Technologies) for 45 min at 37 °C. Tissue from control and experimental groups was digested and stained on the same day to eliminate minor procedure differences. The stromavascular fraction was pelleted by centrifugation at 500g for 10 min, then washed and filtered. Red blood cells are lysed using ACK lysis buffer. Cells were resuspended in 1 ml for counting before staining. For staining, the stromavascular fraction was incubated with FcBlock, surface antibodies for 30 min on ice, in the dark, then washed and stained with Fixable Viability Dye (eBioscience) and intracellular antibodies using cytofix (eBioscience/ThermoFisher). Analysis was performed on a BD LSRII and using FlowJo vX. <u>For sorting B220+:</u> CD3^−^ B220^+^F4/80^−^ CD11b^−^ cells were sorted on a BD FACS Aria into RPMI with antibiotics/antimycotics (ThermoFisher) plus 20% FBS. Cells were pelleted and resuspended in trizol for RNA isolation.

### Lipolysis assay

For *ex vivo* lipolysis assay, adipose tissue was collected and 15 mg was cultured in 100 ul lipolysis buffer (Krebs buffer plus 0.1% glucose and 3.5% fatty acid free BSA; Sigma) in a 96-well plate for 2 h at 37 °C at 450 r.p.m. The glycerol assay (Sigma) was used as per manufacturer’s instructions.

### Western blot

Visceral adipose was snap frozen in liquid nitrogen immediately after harvest. Tissue was homogenized in RIPA buffer containing protease inhibitors. Protein concentration was quantified using the DC protein assay (Bio-Rad) and equal amounts of protein were run on an SDS-PAGE gel and transferred to nitrocellulose membrane. Blots were probed with primary antibodies and then incubated in secondary antibody of the appropriate species (ThermoFisher). Detection occurred using chemiluminescent visualization (Fisher, Bio-rad).

### Whole-mount staining

Staining was performed similar to those previously published. In brief, adipose tissue was collected, fixed in 1% PFA and blocked with 5% BSA. Permeabilization was done using 0.1% Triton-X 100 in 5% BSA. The adipose tissue was then stained in primary antibodies over 1–2 days and secondary antibody, if needed, for 2.5 h in goat blocking serum. Confocal images were acquired using a laser scanning Leica SP8 or Leica SP5 and analyzed using Leica Application Suite AF.

#### For FALC counting

Images were acquired on a Zeiss Inverted Microscope (Axio) or Keyence (BZ-X710) using the stitching function within. FALCs from each tissue were counted using CD3^+^ staining.

#### Histology and pathological analysis

Visceral and subcutaneous adipose tissues were fixed in 10% formalin. Tissues were embedded, sectioned and stained with hematoxylin and eosin or masson’s trichrome by Mouse Research Pathology within Comparative Medicine at Yale University. Pathological analysis of lymphocyte infiltration was accomplished by a double blind procedure without knowledge of the animal’s identity. Two pathologists made independent scores.

### Antibodies

#### For tissue resident labeling

Intravascular labeling was performed by i.v. injection of 2.5ug CD45.2 diluted in 100ul PBS. Mice were euthanized exactly 3 minutes after injection for tissue collection.

#### For flow cytometry analysis

the following antibodies were used: Fixable Viability Dye Aqua; CD3-BV605, B220-PECy7, CD45.2-BV605, CD45.2-FITC, B220-APC, CD45-BV-711, CD4-BV605, FoxP3-APC, Ki67-PECy7, CD19-FITC, B220-AF600, CD8-eF450, CD11b-FITC, MHCII-PECy7, B220-AF700, B220-eF450, CD25-PE, PD1-APC-e780, CD21/35-FITC, CD23-PE, CD23-eF450, F4/80-PE, CD11b-PerCP-Cy5.5, CD3-eF450, B220-FITC, B220-PerCP-Cy5.5.

#### Intracellular staining

Intracellular antigens were detected using the eBioscience Fix/Perm nuclear staining kit as instructed by manufacture-supplied protocol.

#### For whole mount staining

the following antibodies were used: DAPI, B220-APC, B220-PE, CD3-FITC, Prox1 (Abcam; ab101851), Chicken anti-Rabbit-AF488 (Life Tech; A21441)

#### For western blot analysis

the following antibodies were used: pHSL (Ser563; 4139), HSL (4107), ATGL (2439), Beta-Actin (4967) (Cell signaling).

### RNA extraction and gene expression analysis

RNA extraction and purification was performed using the trizol/chloroform method, followed by use of the RNeasy kits (Qiagen) according to manufacturer’s instructions.

### RNA-sequence quality control

Total RNA quality is determined by estimating the 260 nm/*A*280 nm and *A*260 nm/*A*230 nm ratios by nanodrop. RNA integrity is determined by running an Agilent Bioanalyzer gel, which measures the ratio of the ribosomal peaks.

### Library prep

mRNA is purified from approximately 500 ng of total RNA with oligo-dT beads and sheared by incubation at 94 °C. Following first-strand synthesis with random primers, second-strand synthesis is performed with dUTP for generating strand-specific sequencing libraries. The cDNA library is then end-repaired, and A-tailed, adapters are ligated and second-strand digestion is performed by uricil-DNA-glycosylase. Indexed libraries that meet appropriate cutoffs for both are quantified by qRT–PCR using a commercially available kit (KAPA Biosystems). Insert size distribution was determined with the LabChip GX or Agilent Bioanalyzer. Samples with a yield of ≥ 0.5 ng μ l-1 are used for sequencing.

### Flow cell preparation and sequencing

Sample concentrations are normalized to 10 nM and loaded onto Illumina High-output flow cells at a concentration that yields 150–250 million passing filter clusters per lane. Samples are sequenced using 75 bp single-end sequencing on an Illumina HiSeq 2000 according to Illumina protocols. Each sample has a 6-bp index that is incorporated during the library prep. The index sequence is read using a different sequencing primer than the 75-bp sequencing read. The index read happens immediately after the 75-bp read. Data generated during sequencing runs are simultaneously transferred to the YCGA high-performance computing cluster. A positive control (prepared bacteriophage Phi X library) provided by Illumina is spiked into every lane at a concentration of 0.3% to monitor sequencing quality in real time.

### Data analysis and storage

Signal intensities are converted to individual base calls during a run using the system’s real time analysis (RTA) software. Sample de-multiplexing was performed using Illumina’s CASAVA 1.8.2 software suite. The data are returned to the user if the sample error rate is less than 2% and the distribution of reads per sample in a lane is within reasonable tolerance.

### Primary data analysis of RNA-Seq data

Raw fastq-files were mapped to the murine genome version mm10 using HISAT version 0.1.7-β (Kim et al., 2015) using the default options. The resulting BAM files were imported into Partek® Genomics Suite® software, version 6.6 Copyright©; 2017 (PGS) and reads were quantified using the mm10-RefSeq Transcripts database version 2016-02-02 to obtain read counts for each individual RefSeq gene.

Next, the raw read counts were imported into R and further analysis have been carried out using the R package DEseq2 (Love et al., 2014). Following, normalization we determined differentially expressed genes using DESeq2 with standard settings and IHW (Ignatiadis et al., 2016) filtering. Genes were considered as differentially expressed (DE) if the adjusted *p* value (IHW, weighted Benjamini and Hochberg correction) was less than 0.1. The differences in gene expression were visualized as the dependency between the negative decadic logarithm of the p-value and the difference in gene expression as log2 fold change (volcano plot).

### B-Cell macrophage interaction model

To create a transcriptome-informed model of possible cell-to-cell ligand-receptor interactions we utilized the Fantom5 receptor-ligand database (Ramilowski et al., 2015). First, we downloaded the receptor-ligand interactions [http://fantom.gsc.riken.jp/data/, October/2016] and replaced the human receptor/ ligands with murine orthologs using ENSEMBL biomart (Ensembl Genes 92, GRCh38.p12). Genes that lacked a murine ortholog in this database were manually curated using the NCBI gene database. Interactions were filtered by DE genes of the respective comparison and visualized using the R package igraph 1.2.1 [https://cran.r-project.org/web/packages/igraph/index.html].

## QUANTIFICATION AND STATISTICAL ANALYSIS

### Experimental design

Blinding of investigators was not possible during experiments. Control and experimental groups were randomly assigned by cage. All experiments contained littermates and non-littermates, which were both randomly assigned to control and experimental group. Young control mice were raised or obtained from Jackson Laboratory (wild-type; C57BL6/J). Statistical significance was calculated by a two-tailed Student’s *t*-test or ANOVA using a post-hoc test to correct for multiple hypotheses. \**P* < 0.05; \*\**P* < 0.005; \*\*\**P* < 0.001; \*\*\*\**P* < 0.0001. GraphPad Prism was used to define statistical outliers, which were then excluded from data analysis. A confidence interval of 95% was used for all statistical tests. All data were assumed to be normally distributed, unless the standard deviation was identified as significantly different between groups. All statistical tests were performed using GraphPad Prism v7 for Windows (GraphPad Software). Data are expressed as mean ± s.e.m. Biological replicates and the number of independent experiment repetition are described in the figure legends.

## DATA AND SOFTWARE AVAILABILITY

The RNA sequencing data will be uploaded to publically available database upon acceptance of this manuscript. Previously published datasets that were used in the analysis are indicated by GEO number in the text.

**Supplemental 1. Related to Figure 1.**

a. Gating strategy to identify tissue residents using flow cytometry

b. Quantification of CD19^+^ cells in the visceral adipose tissue of 2 and 24 month old male or female mice that were bred and aged at Pennington Biomedical Research Center.

c. Contour plots (gated through CD45^+^ live cells) of B220^+^ resident cells in VAT, SAT and BAT.

d. Whole mount fluorescent imaging of mesenteric and gonadal visceral adipose depots from female mice at 7 or 24 months of age. CD3 (red), B220 (blue)

e. Whole mount confocal imaging of FALCs in VAT of 22month old wild-type mouse. Top: CD3 (green) and B220 (red). Bottom: CD4 (red), B220 (white), and DAPI (blue).

f. CD11b MFI on B220^+^ cells from visceral adipose tissue of female or male mice at 3 or 24 months of age. Each symbol represents one mouse

g. Quantification of the percentage of CD11b^+^ and CD11b^−^ out of the total B220^+^ residents in VAT from 3 or 24 month old male and female mice.

h. Quantification of B cell subsets in spleen, MAT and VAT from 24 month old female mice.

i. Contour plots (gated through CD45^+^ B220^+^ live cells) of CD21 and CD23 expression on B cells from 3 month old mice after no stimulation (UNTX) or stimulation with adipose media from 3 or 24 month old mice.

j. Whole mount confocal imaging of B220^+^ cells (green) in lymphatic vessel of visceral adipose tissue from 22 month old ProxTom mice.

k. Whole mount confocal imaging of B220+ mesenteric lymph nodes (red) in 22 month old wild-type mice stained with Prox1 antibody (green).

l. Whole mount fluorescence imaging of B220^+^ FALCs (red) in 22 month old wild-type mice stained with Prox1 antibody (green).

m. Whole mount confocal imaging of B220^+^ FALCs (red) in 22 month old wild-type mice stained with Prox1 antibody (green). Two representative images are shown.

All error bars represent mean±SEM. *<0.05, **<0.01, ***<0.005. Statistical significance was determined by an ANOVA with posthoc test to adjust for multiple corrections.

**Supplemental 2. Related to Figure 2.**

a. Single color images for the whole mount confocal imaging of FALC in VAT of 22month old mTmG;LysMcre with B220 (yellow) and DAPI (blue) antibody staining. mTomato expressed on all cells and mGFP on myeloid cells.

b. Quantification of CD4^+^ and CD8^+^ T cells (as a percentage of total SVF) in VAT from 3 or 24 month old WT and Nlrp3-/- mice

c. Quantification of CD11b^+^ cells, as a percentage of CD45^+^ live cells, in WT and Nlrp3-/- mice.

d. Quantification of CD4^+^ T cells as a percentage of CD45^+^ live cells, in WT or Nlrp3-/- mice

e. Quantification of B220^+^ cells as a percentage of CD45^+^ live cells, in WT and Nlrp3-/- mice.

f. Mean gene expression of chemokines from 3 month old WT, 24 month WT and Nlrp3-/- adipose macrophages. All chemokines containing ccl or cxcl within genes were selected from previously performed RNA sequencing analysis of adipose tissue macrophages. Genes were ordered from highest to lowest according to expression level in 24 month old WT macrophages and hierarchical clustering analysis was performed.

g. Schematic showing the interaction model between macrophages and B-cells. Displayed macrophage interaction partners are filtered by their differential expression between wild-type and Nlrp3-deficient macrophages and visualized in a network with their corresponding interaction partners. An edge shows an interaction listed in the FANTOM5 interaction database. Interactors are color-coded by log2FC, if they are significantly regulated between wild-type and Nlrp3-deficient macrophages or B-cells, respectively.

All error bars represent mean±SEM. *<0.05, **<0.01, ***<0.005. Statistical significance was determined by an ANOVA with posthoc test to adjust for multiple corrections.

**Supplemental 3. Related to Figure 3.**

a. Body-weight, visceral adipose tissue weight and visceral adipose tissue cellularity in WT or GHRKO mice at 20 months of age.

All error bars represent mean±SEM. *<0.05, **<0.01, ***<0.005. Statistical significance was determine by an ANOVA with posthoc test to adjust for multiple corrections.

**Supplemental 4. Related to Figure 4.**

a. Schematic to describe intra-adipose depletion.

b. Body-weight in mice at intra-adipose injection at time points indicated until euthanasia and tissue harvest. Right graph quantifies final body-weight of individual mice. Each symbol represents one animal.

c. VAT weight of mice treated as described.

d. Histogram plots showing B220^+^ cells in tissues from 20 month old mice given intra-adipose injection of ISO or CD20 mAb. Values represent the mean percentage of B220^+^ cells.

e. Schematic to describe systemic depletion.

f. Quantification of body-weight in mice treated as described. Each symbol represents an individual mouse.

g. Quantification of visceral adipose tissue weights in mice that were aged and treated as described.

h. Representative contour plots gated through live CD45^+^ CD4^+^ cells, showing CD25 and FoxP3 fluorescent expression in spleen or visceral adipose tissue of mice as described. Values represent mean for that condition.

i. Histogram plots showing PD1 expression on live CD45^+^ CD4^+^ T cells from spleen or visceral adipose tissue of mice as described. Values represent the mean percent for each condition.

j. Insulin tolerance test in 21 month old mice that were treated as described.

k. Area under the curve (AUC) of mice challenge with insulin.

l. Quantification of glycerol levels in the supernatant of visceral adipose tissue from mice that were aged and treated as labeled, prior to harvest and stimulated with 2uM isopreteronol to induced lipolysis.

m. Rectal temperature at day 0 and day 3 in 3-8 month old mice and 20-25 month wild-type mice exposed to four degrees.

All error bars represent mean±SEM. *<0.05, **<0.01, ***<0.005. Statistical significance was determined by an ANOVA with posthoc test to adjust for multiple corrections.

