## Supplemental figure 1-4 for "Aging induces Nlrp3 inflammasome dependent adipose B cell expansion to impair metabolic homeostasis"

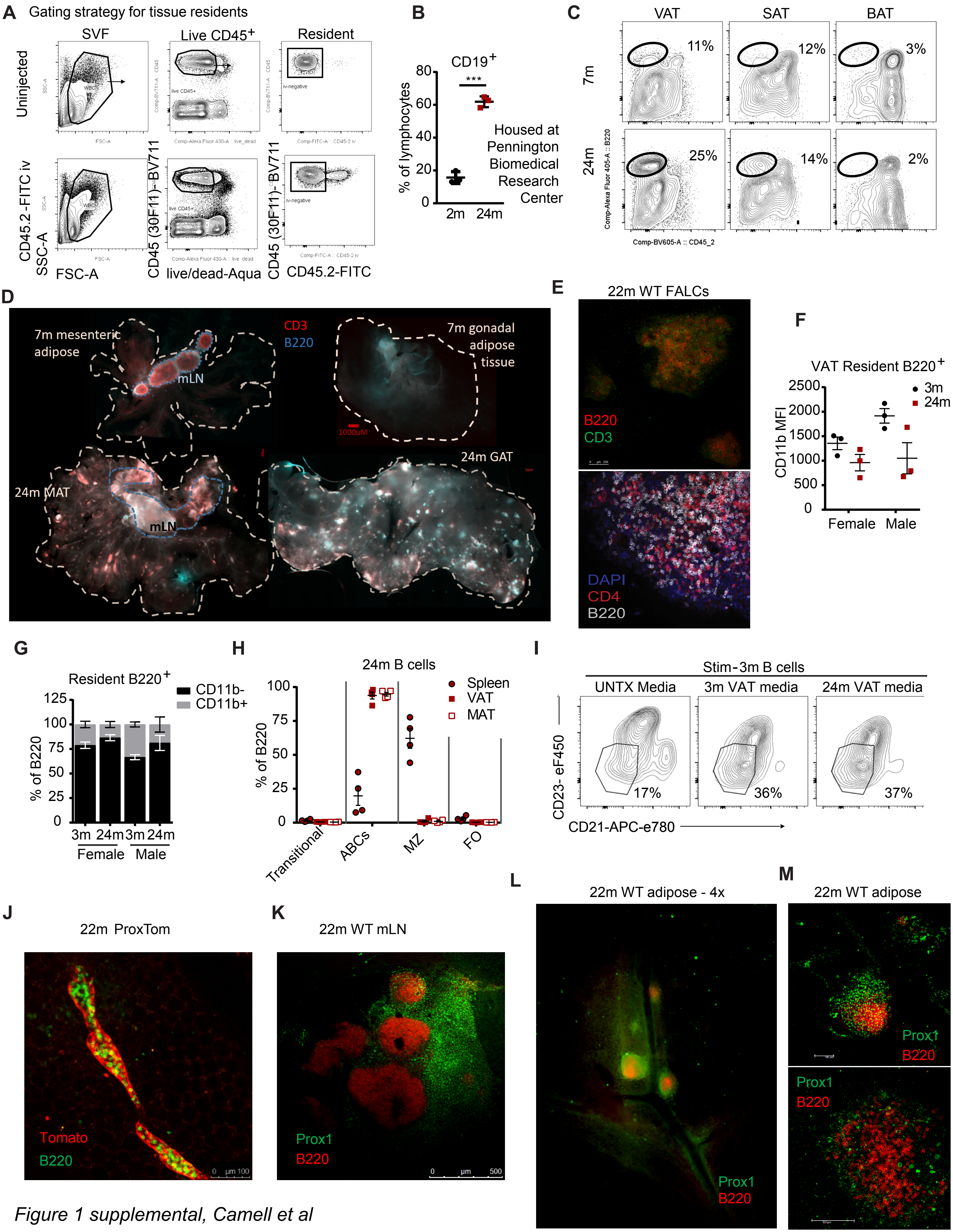


*Figure 1 supplemental, Camell et al*

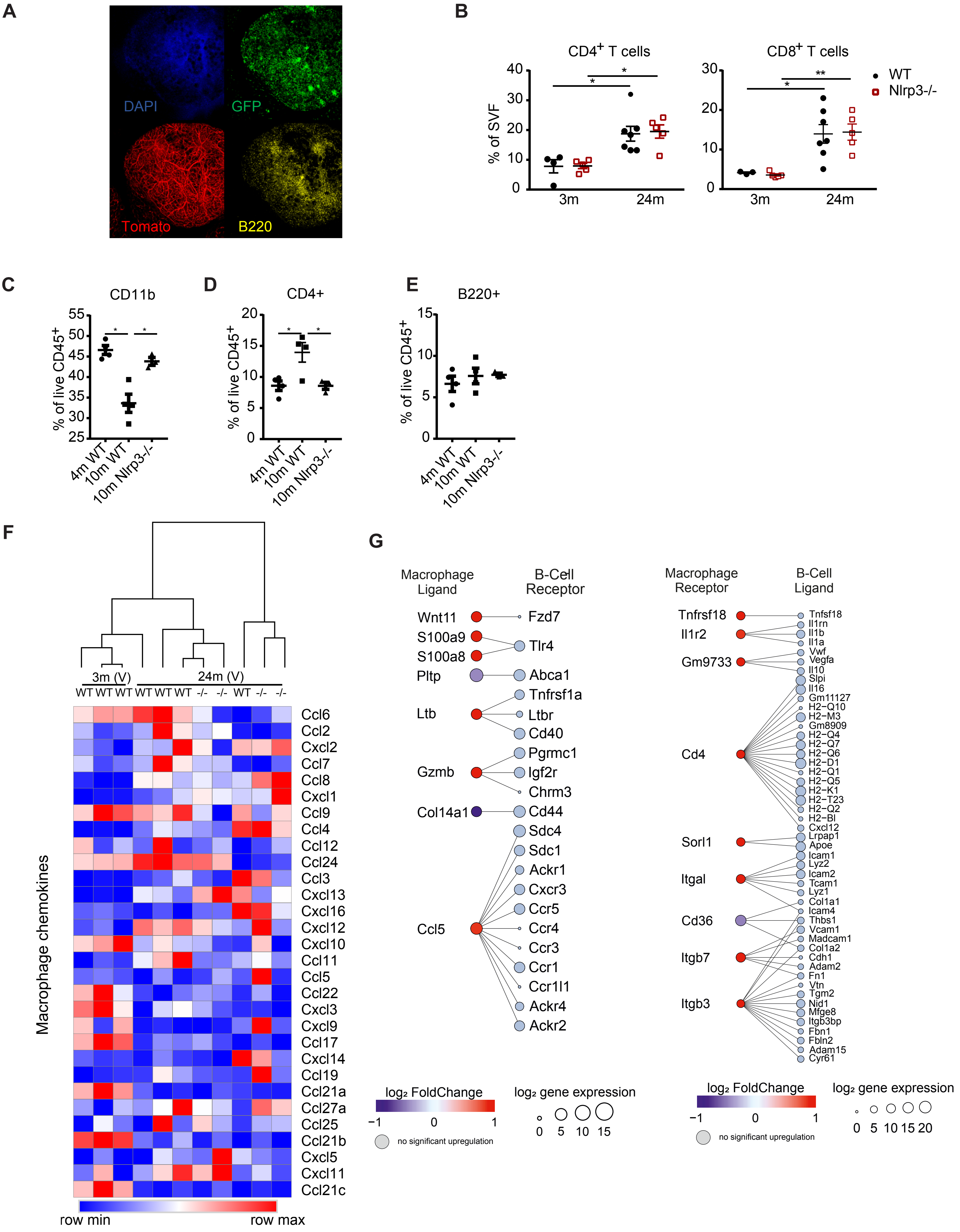


*Supplemental figure 2, Camell et al*

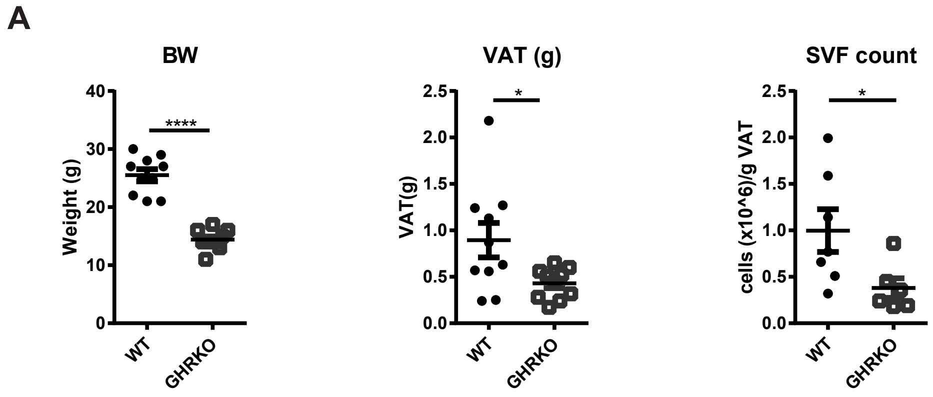


*Supplement figure 3, Camell et al*

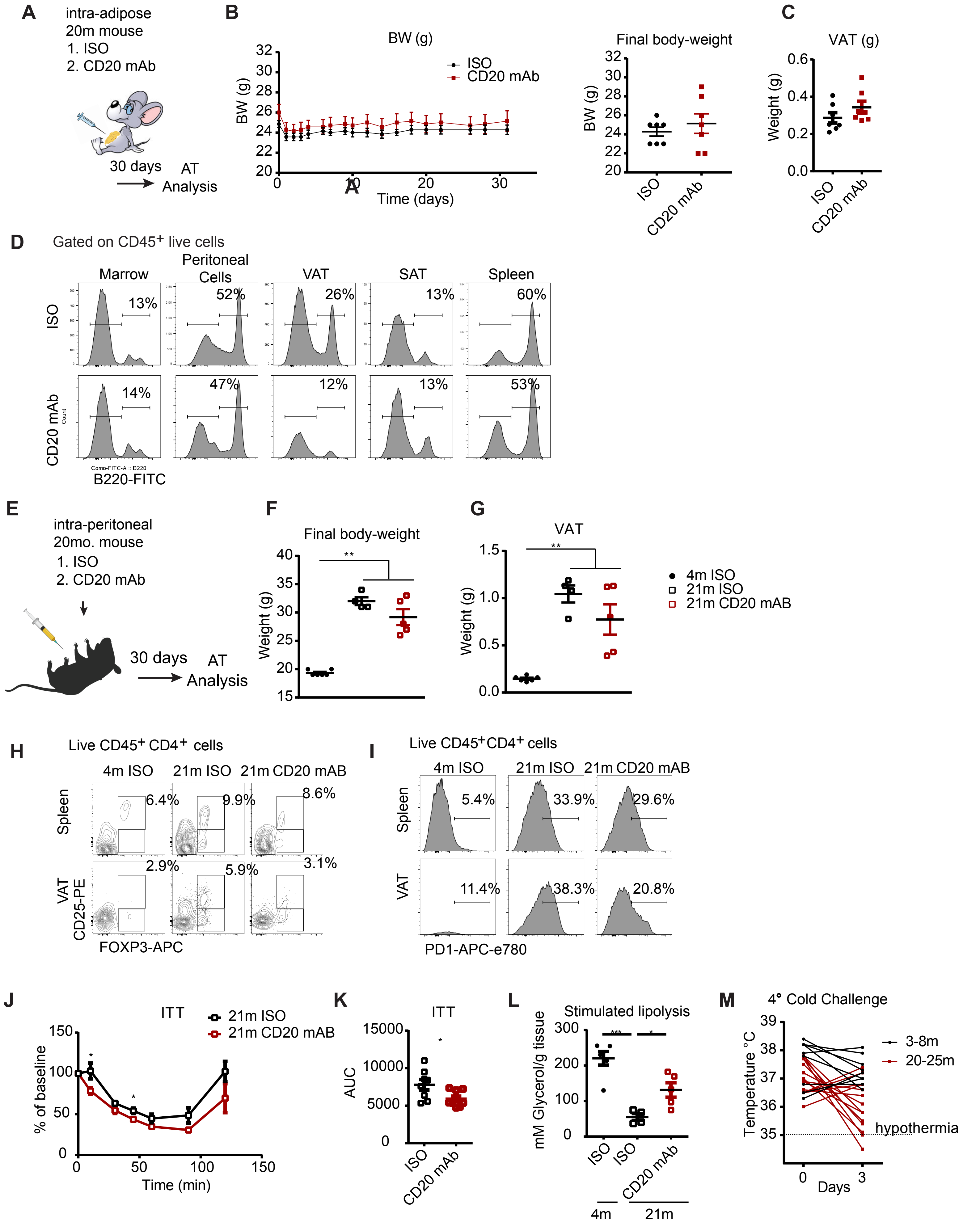


*Supplemental Figure 4, Camell et al*
